# CDK11 activates CDK12 to trigger the elongation of RNA Polymerase II

**DOI:** 10.1101/2025.09.27.678947

**Authors:** Haolin Liu, Lingbo Li, Junfeng Gao, Sonia Leach, Pengcheng Wei, Qianqian Zhang, Hua Huang, Thibault Houles, Philippe P. Roux, Gongyi Zhang

## Abstract

CDK11 is an essential cyclin-dependent kinase in higher eukaryotes, yet its specific substrates and functional roles have remained elusive. Our findings reveal that CDK11 is functionally analogous to the yeast Bur1 kinase, phosphorylating the repeat region of hSpt5 and the linker domain of Rpb1 in RNA Polymerase II (Pol II). Inhibition of CDK11 results in a significant reduction of active Pol II not only at transcription start sites but also along gene bodies. Further investigation finds that CDK11 is crucial for the phosphorylation and activation of CDK12, which is essential for the elongation of Pol II. Moreover, we find that CDK10 can partially compensate for the function of CDK11. Combining with earlier functional elucidation of CDK9 from us, the consequent roles of CDK7/8, CDK9, CDK10/11, and CDK12/13 in transcription regulation in metazoans are established.

## Introduction

Promoter-proximal RNA Polymerase II (Pol II) pausing regulates the expression of approximately 30-60% of genes in higher eukaryotes (Jonkers and Lis 2015), which is likely regulated by the presence of arginine-methylated nucleosomes at the +1 position from transcription start sites (Weber, Ramachandran et al. 2014, Liu, Ramachandran et al. 2020, Liu, Wei et al. 2022). Our research has identified that specific members of the Jumonji protein family, JMJD5 and JMJD7, possess the dual endo- and exo-peptidase activities necessary to cleave histone tails containing methylated arginines, thereby generating fragile “tailless nucleosomes” in higher eukaryotes (Liu, Wang et al. 2017, Liu, Wang et al. 2018). Detailed analyses revealed that JMJD5 specifically targets +1 nucleosome across a broad set of genes (Liu, Ramachandran et al. 2020) and functions in concert with CDK9 to release paused Pol II into active transcription elongation (Liu, Ramachandran et al. 2020). CDK9 is recruited to Pol II by BRD4 and JMJD6 following its release from the 7SK snRNP complex (Hong, Zang et al. 2010, Lee, Liu et al. 2020). We propose that promoter-proximal Pol II pausing is intricately linked with multiple processes, including the high turnover of histones in non-proliferating cells, CDK9 activity, the unique pattern of phosphorylation of the Pol II C-terminal domain (CTD) by CDK9, the recruitment of CDK9 to Pol II by BRD4 and JMJD6, and potentially p300/CBP (Lee, Liu et al. 2020, Liu, Ramachandran et al. 2020, Zhang 2024).

Studies have shown that the yeast kinase, Bur1, specifically phosphorylates the C-terminal repeats region of Spt5 (Zhou, Kuo et al. 2009, Qiu, Hu et al. 2012, Wier, Mayekar et al. 2013) and the linker region of heptad repeats in the C-terminal domain and the main body of Rpb1 of Pol II. These events are responsible for the recruitment of elongation complex Paf1c (Mayekar, Gardner et al. 2013) and Spt6 (Sdano, Fulcher et al. 2017, Chun, Joo et al. 2019), respectively. However, inhibition of Bur1 does not affect the phosphorylation status of the CTD of Pol II (Booth, Parua et al. 2018, Chun, Joo et al. 2019), contradicting the prevailing view that Bur1 is the ortholog of CDK9 in metazoans (Eick and Geyer 2013). This suggests that yeast Bur1 (Prelich and Winston 1993) has a functionally unsuspected analog in higher eukaryotes.

Cyclin-dependent kinases (CDKs) not only participate in the regulation of the cell cycle but also in transcription initiation and elongation in eukaryotes. Due to their critical roles, most of these proteins are conserved from simple eukaryotes (yeast) to higher eukaryotes (humans). However, CDK11 remains an enigmatic candidate in the animal kingdom, as its ortholog in yeast has not been identified (Malumbres 2014). CDK11 was cloned three decades ago (Bunnell, Heath et al. 1990) and characterized in the late 1990s and early 2000s (Loyer, Trembley et al. 1998, Trembley, Hu et al. 2002, Hu, Mayeda et al. 2003, Trembley, Hu et al. 2003, Li, Inoue et al. 2004, Loyer and Trembley 2020) by pioneering work from the late Dr. Vincent Kidd’s group. CDK11 knockout mice die at an early pre-implantation stage (E3.5-E4.5) (Li, Inoue et al. 2004), earlier than the organogenesis defects seen in CDK9 knockouts (∼E9.5) (Dickinson, Flenniken et al. 2016), highlighting CDK11’s critical role in early embryonic development.

A critical question remains regarding the functions of CDK11. CDK11 has been shown to phosphorylate several splicing factors, thus controlling pre-mRNA splicing (Loyer, Trembley et al. 1998, Hu, Mayeda et al. 2003). Recent findings identify SF3B1 as a target of CDK11 (Hluchy, Gajduskova et al. 2022), which further supports the observed role of CDK11 in the regulation of splicing (Loyer, Trembley et al. 1998, Hu, Mayeda et al. 2003). However, CDK11 may have additional functions, including, for example, direct or indirect effects on Pol II. Thus CDK11 inhibition has been reported to block transcription from promoters with both TATA boxes and GC-rich sequences (Trembley, Hu et al. 2002), suggesting that CDK11 participates in the regulation of both SAGA-dependent and TFIID-dependent transcription mechanisms that govern all Pol II-transcribed genes (Rossi, Kuntala et al. 2021). The recent discovery that the specific CDK11 inhibitor, OTS964, affects the levels of nearly all transcripts (Lin, Giuliano et al. 2019, Hluchy, Gajduskova et al. 2022) further corroborates earlier findings (Trembley, Hu et al. 2002). In this report, we find that CDK11 phosphorylates the C-terminal repeat regions of hSpt5.

This phosphorylation is essential for recruiting Pol II elongation factors such as the Paf1 complex (Paf1c) (Zhou, Kuo et al. 2009, Qiu, Hu et al. 2012, Mayekar, Gardner et al. 2013, Wier, Mayekar et al. 2013). CDK11 also phosphorylates the linker region of the C-terminal heptad repeats and the main body of Rpb1, the largest subunit of Pol II. This phosphorylation is critical for the recruitment of the essential elongation factor, Spt6 (Sdano, Fulcher et al. 2017, Chun, Joo et al. 2019). However, we observed no significant changes in the phosphorylation status of Ser5-CTD of Pol II after treatment with OTS964, although a dramatic reduction in Ser2-CTD phosphorylation was noted. Consistent with previous reports showing that CDK11 does not directly phosphorylate the heptad repeats of CTD Pol II (Trembley, Hu et al. 2003). To our surprise, we find that CDK11 and CDK10 are vital for phosphorylation and activation of CDK12 to trigger the phosphorylation of Ser2-CTD of Pol II (Ser2ph-CTD of Pol II), which is essential for the subsequent elongation of Pol II. We also find that CDK11/10 may work on other factors, such as termination factors, besides splicing factors, along gene bodies.

### CDK11 phosphorylates the C-terminal repeats of human Spt5 (hSpt5)

From accumulating data generated by us and others, it is clear that CDK9 from higher eukaryotes is not the ortholog of yeast Bur1. To investigate the potential ortholog of Bur1 in humans or mice, the protein sequence of yeast Bur1 was uploaded onto the BLAST program in search of potential homologs either in the human genome or the mouse genome. It is quite striking that both CDK11A and CDK11B, with a signature motif of “PITSLREA/B”, show up as top candidates with CDK9 followed (**Fig. S1**), suggesting that functionally uncharacterized CDK11 is a potential ortholog of yeast Bur1. Based on the previous functional characterization of Bur1, which is specific to phosphorylating the C-terminal repeat (CTR) of Spt5 (Zhou, Kuo et al. 2009, Qiu, Hu et al. 2012, Wier, Mayekar et al. 2013), we reason that CDK11 should carry out a similar role on human hSpt5. Interestingly, the sequence of the CTR of human hSpt5 is quite different from Saccharomyces cerevisiae Spt5 but almost identical to that of Schizosaccharomyces pombe (**Fig. S2**). To read out the activity of CDK11 on hSpt5, we raised a rabbit polyclonal antibody (pAb) against a phosphorylated CTR sequence of hSpt5 characterized earlier (**Fig. 1a**). Our rabbit pAb specifically recognizes hSpt5 (Lane 1, **Fig. 1b**), which is cross-confirmed by a monoclonal antibody against hSpt5 with an unphosphorylated epitope (Lane 1, **Fig. 1b**). To verify if the rabbit pAb is specific for phosphorylated CTR generated by CDK11, the inhibitor of CDK11, OTS964, was applied to the growing HEK293T cell lines in a gradient format. As predicted, OTS964 reduced the phosphorylation status of hSpt5 (**Fig. 1b**), proving that the phosphorylation of CTR of hSpt5 is generated by CDK11 and the rabbit pAb raised could read out the activity of CDK11. Finally, a CTR-hSpt5-MBP fusion protein was used to test the activity of CDK11 *in vitro*. Again, CDK11 can phosphorylate CTR of hSpt5 while OTS964 inhibits the activity of CDK11 (**Fig. 1c**). These phosphorylation assays demonstrate that hSpt5 is a cognate substrate of CDK11.

**Figure 1.**
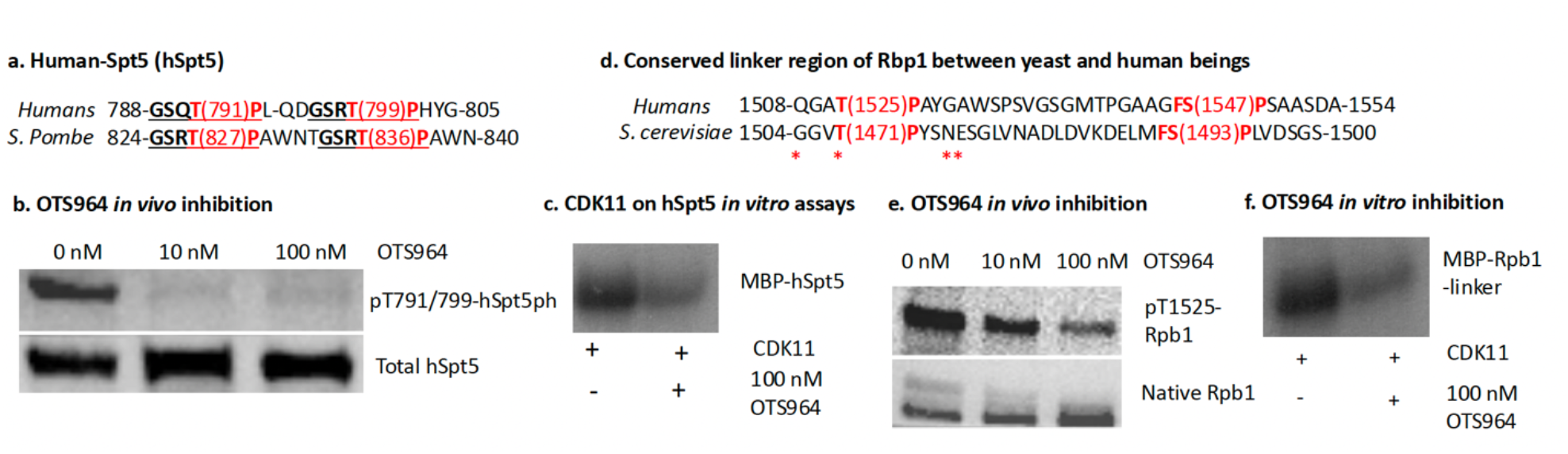
hspt5 and the linker of Rpbl are cognate substrates of CDK11. **a**. A two-repeat of human hSpt5 from 786-804 with two phosphor groups at Thr791 and Thr799 was used to raise rabbit polyclonal antibodies. **b**. OTS964 inhibition leads to the disappearance of phosphorylated hSpt5 (hspt5ph) *in viva in HEK2937 cells*. **c**. In vitro kinase assay showed OT5964 inhibits phosphorylation of a fusion fragment of Spt5 and MBP by CDKIL. **d**. sequence alignment of linker region of Rop1 of yeest and human. **e**. COK11 phosphorylates the linker of Rpbl and is inhibited by OTS964 *in vivo* in HEK293T cells. **f**. *In vitro* assay of CDK11 on the linker of Rpbl with and without CT5964.

### CDK11 phosphorylates the linker region of heptad repeats of CTD and the body of Rpb1 of Pol II

A study showed that phosphorylation of a region that lies between the heptad repeats and the main body of Rpb1 of Pol II is required to recruit Spt6 (Sdano, Fulcher et al. 2017). A later report showed that, in yeast, Bur1 is responsible for this phosphorylation (Chun, Joo et al. 2019). The sequences of the linker region of Rpb1 in yeast and humans are quite similar (**Fig. 1d**) and, indeed, we found that a rabbit polyclonal antibody (anti-pT1471 raised against a phosphorylated Rpb1 linker from yeast (Sdano, Fulcher et al. 2017) also recognized the corresponding phosphorylated site of human Rpb1. Therefore, to find out if CDK11 was responsible for the phosphorylation of the human Rpb1 linker, we examined whether the CDK11 inhibitor, OTS964, would inhibit the reaction in vivo and in vitro. OTS964 did have this inhibitory activity (**Fig. 1e**), although not as dramatically as its effects on hSpt5ph (**Fig. 1b**).

Again, the Rpb1-linker-MBP fusion protein was expressed and purified for in vitro kinase assays, the activity of CDK11 toward the liker of Rpb1 was inhibited by OTS964 (**Fig. 1f**). These data showed that CDK11 has activities that are similar to Bur1 in that can phosphorylate the linker region within Rpb1.

### Inhibition of CDK11 does not affect the levels of Ser5ph-CTD of Pol II but leads to a dramatic drop in Ser2ph-CTD of Pol II

Both Spt5 and Spt6 are essential in yeast, suggesting that they play pivotal roles, via Bur1, in the regulation of activities of Pol II. Both Spt5 and SPT6 are essential in animals, too. However, the kinase(s) responsible for the phosphorylation of Spt5 and recruitment of Spt6 remain unknown. We suggest that CDK11 may have the same role as Bur1 since knockout of CDK11 in mice is embryonic lethal, mice die at E3.5 to E4.5 (Li, Inoue et al. 2004), an earlier stage than mice lacking CDK9 die at E8 and E9 (Dickinson, Flenniken et al. 2016). To find out whether CDK11 participates in the regulation of Pol II, we investigated the inhibition of CDK11 by OTS964 to check if it affects the modification of the CTD of Pol II. Inhibition of CDK11 did not affect the level of Ser5ph-CTD of Pol II (**Fig. 2a**) as readout by the antibody against Ser5ph-CTD, 3E8 (Chapman, Heidemann et al. 2007). However, inhibition of CDK11 led to a dramatic drop of Ser2ph-CTD of Pol II as detected by 3E10 (Chapman, Heidemann et al. 2007) (**Fig. 2b**). To confirm this observation, a time course CUT&TAG-seq of elongating Pol II (active Pol II with Ser2ph-CTD of Pol II) was carried out. As we predicted, Ser2ph-CTD of Pol II started to drop at the transcription start sites (TSS) of genes after 2.5 minutes of inhibition of CDK11 by OTS964 (**Fig. 2c**). The effect continued from 5 minutes and 10 minutes for all genes with defined and no-overlapping (total of 4,472) transcription start sites (**Fig. 2c**). This was also true for active Pol IIs within the gene bodies, which revealed a gradual disappearance of Ser2ph-CTD of Pol II with time (**Fig. 2d**). These results suggest that inhibition of CDK11 shuts down all Pol II transcribed genes in HEK293T cells, which may explain why CDK11 is essential role for the early embryonic development of animals as reported early (Li, Inoue et al. 2004).

**Figure 2.**
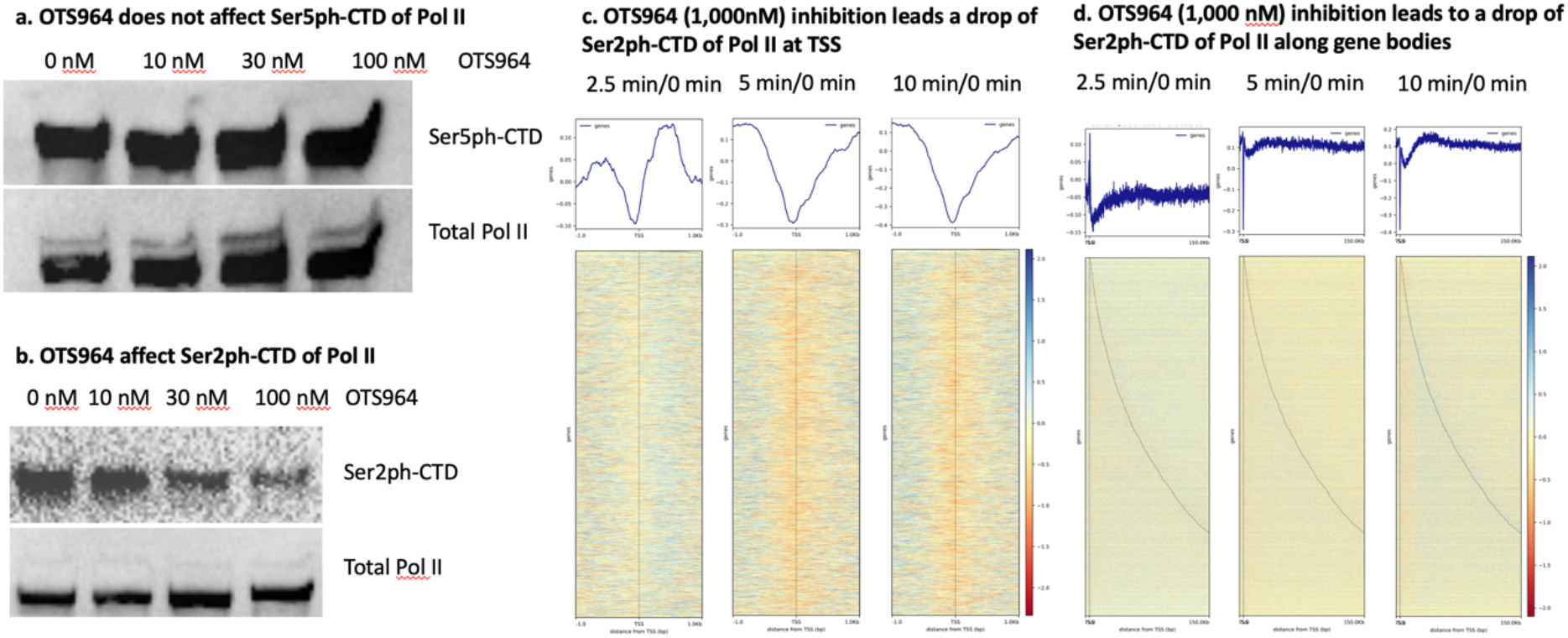
OTS964 inhibition leads to a drop of active Pol IIs (Ser2ph-CTD of Pol II). **a**. OTS964 does not affect SerSph-CTD of Pol II. **b**. OTS964 leads to a drop of Ser2ph-CTD of Pol II. **c**. OTS964 (1,000 nM) leads to a dramatic drop of Ser2ph-CTD of Pol II at transcription start sites (TSS) with time course. **d**. OTS964 (1,000 nM) leads to a dramatic drop of Ser2ph-CTD of Pol II along gene bodies with time course.

### CDK12 is a cognate substrate of CDK11

It was reported that CTD of Pol II is not a cognate substrate of Bur1 in yeast (Booth, Parua et al. 2018, Chun, Joo et al. 2019) or CDK11 from mice (Trembley, Hu et al. 2003). However, our data showed that the level of Ser2ph-CTD of Pol II dramatically drops after inhibition of CDK11 (**Fig. 2b, 2c, and 2d**). It is of great interest to address the underlying mechanism of this phenomenon. It was well established that Ser2ph-CTD of Pol II is generated by Ctk1 in yeast and CDK12/13 in animals (Bartkowiak, Liu et al. 2010). We reasoned that CDK11 may target CDK12/13 and thus indirectly affect the phosphorylation of Ser2-CTD of Pol II. Interestingly, a recent report from one of the authors’ groups showed that phosphorylation of T548 of CDK12 is essential for the activities of CDK12 in human cancer cell lines (Houles, Lavoie et al. 2022). We suspect that CDK12 could be a cognate substrate of CDK11. As predicted, inhibition of CDK11 by OTS964 led to a dramatic drop in the levels of active CDK12 (pT548-CDK12) (**Fig. 3a**). To find out if this is also true for individual genes, we used CUT&TAG-seq against native CDK12 (nCDK12) and active CDK12 (pT548-CDK12), respectively. Inhibition of CDK11 led to the accumulation of the native form of CDK12 at transcription start sites genome-wide (**Fig. 3b**). However, the active form of CDK12 started to drop with the inhibition of CDK11 (**Fig. 3c**). This result supports our idea that CDK11 phosphorylates and activates CDK12 to subsequently generate elongating active Pol IIs (Ser2ph-CTD of Pol II).

**Figure 3.**
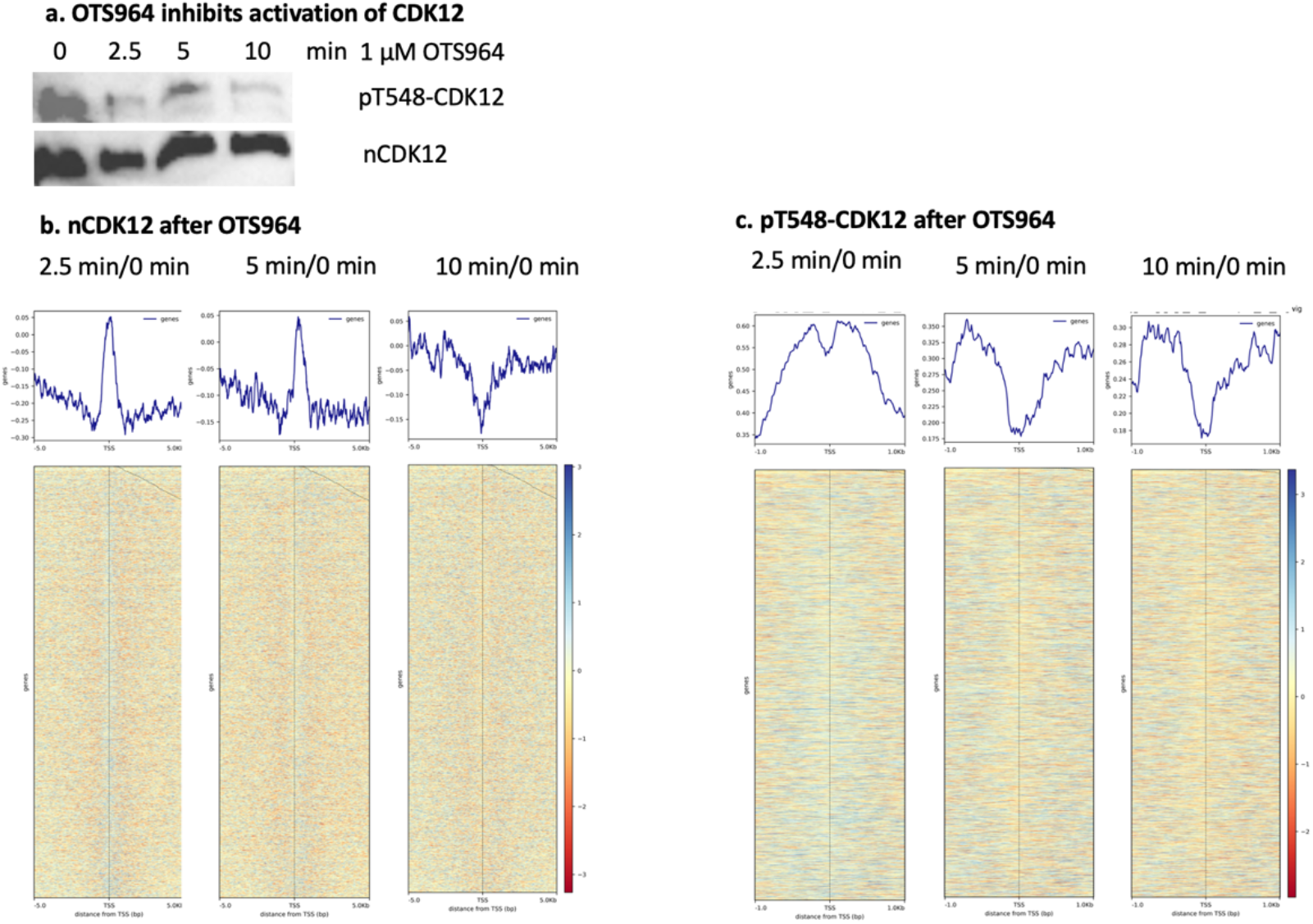
OTS964 inhibition leads to an accumulation of native CDK12 but a decrease of active CDK12 at TSS. **a**. OTS964 inhibition leads to a decrease of active CDK12. **b**. Native CDK12 accumulates at TSS. **c**. active CDK12 decreases at TSS after OTS964 inhibition at TSS.

### CDK10 partially compensates for the function of CDK11

As we demonstrated, inhibition of CDK11 by OTS964 led to a dramatic drop in the levels of pT791 of hSpt5 (**Fig. 1b**), but the effects are milder toward pT1525 of Rpb1 (**Fig. 1e**) and pT548 of CDK12 (**Fig. 3a**). We wondered if CDK11 was the only kinase to carry out the phosphorylation of these characterized substrates. In this regard, shRNAs of CDK11 were used to knock down CDK11, and the phosphorylation levels of these substrates were monitored. Five shRNAs of CDK11 could successfully knock down CDK11 in HEK293T cells (**Fig. 4a**). The knockdown of CDK11 also leads to down-regulation of pT791 of hSpt5 (**Fig. 4b**), suggesting CDK11 is responsible for the phosphorylation of hSpt5. However, we found that the knockdown of CDK11 alone did not lead to a fall in the level of pT1525 of the linker of Rpb1 (**Fig. S3**) or a fall the level of pT548 of CDK12 (the second row, **Fig. 4d**). The results suggest that other kinase (s) which also function on both Rpb1 and CDK12 could be targeted by OTS964. In mice and humans, CDK10 was found to be highly similar to CDK11 (Malumbres 2014). This is true when CDK10 and CDK11 are aligned by sequence (**Fig. 4S**). Recent reports showed that OTS964 also specifically inhibits CDK10 though with a much higher IC50 of ∼5.3 μM (Duster, Ji et al. 2022) than that of CDK11 with Kd of ∼40 nM (Duster, Ji et al. 2022). It suggests that in the case of knockdown of CDK11 alone, CDK10 could compensate for the function of CDK11. To confirm this, we applied a CDK10 inhibitor, Dinaciclib with IC50 of ∼ 1 μM (∼916 nM) (Robert, Johnson et al. 2020, Duster, Ji et al. 2022), to the medium of HEK293T cells coupled with the shRNA knockdown of CDK11. 1 μM Dinacilib led to the drop of both p1525-Rpb1 (**Fig. 4c**) and pT548-CDK12 (top row, **Fig. 4d**). The results suggest that CDK10 could act as a redundant homolog to CDK11 to phosphorylate both Rpb1 and CDK12.

**Figure 4.**
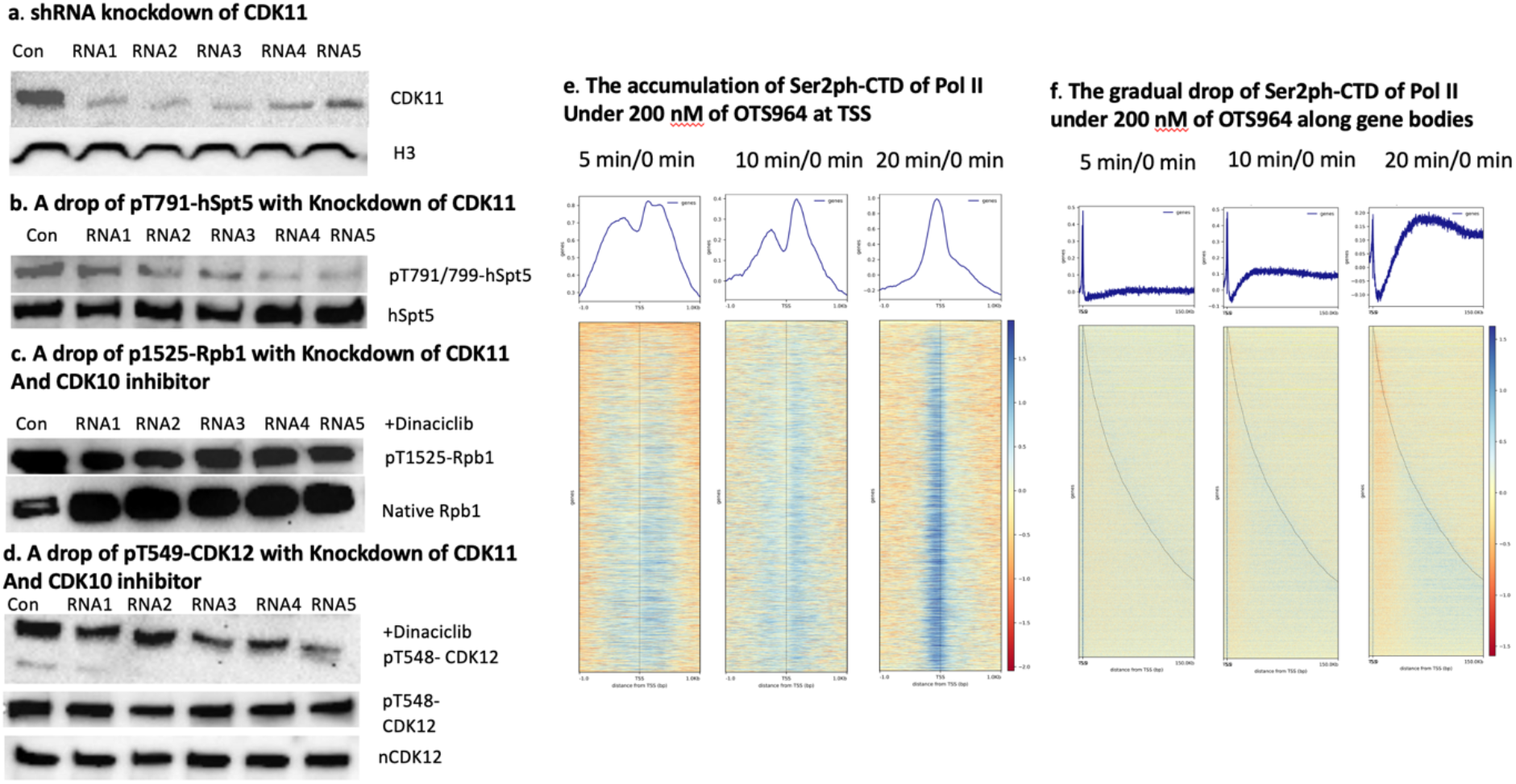
CDK1O is a homolog of CDK11 working on Rpb1 and CDK12. **a**. shRNA knockdown of CDK11 in HEK293T cells. **b**. A drop of pT791-hSptS with only a knockdown of CDK11. **c**. pT1525-Rpbl drops with knockdown of CDK11 and inhibition of CDK1O by Dinaciclib. **d**. pT548 drops with knockdown of CDK11 and inhibition of CDK1O by Dinaciclib. **e**. The accumulation of Ser2ph-CTD of Pol II Under 200 nM of OTS964 at TSS. **f**. The gradual drop of Ser2ph-CTD of Pol II under 200 nM of OTS964 along gene bodies.

An early report showed that inhibition by OTS964 led to the accumulation of Ser2ph-CTD of Pol II at TSS (Hluchy, Gajduskova et al. 2022). The results are not consistent with our data (**Fig. 2**). However, 30 nM OTS964 was used for the ChIP-seq analysis in the published report (Hluchy, Gajduskova et al. 2022), a concentration that is much lower than the IC_50_ of CDK10. In our experiments involving CUT&TAG-seq data of Ser2ph-CTD of Pol II and CDK12, we used 1000 nM of OTS964, although we used lower concentrations of OTS964 for the western blot analyses (**Fig. 2 and Fig. 3**). Thus the differences in results in reference (Hluchy, Gajduskova et al. 2022) by comparison with our own data may be due to our use of higher concentration of the inhibitor. To check this, we carried out CUT&TAG-seq on Ser2ph-CTD of Pol II under 200 nM OTS964 and found that the lower concentration of OTS964 did lead to the accumulation of Ser2ph-CTD of Pol II at transcription start sites (**Fig. 4e**). However, the level of Ser2ph-CTD gradually dropped along gene bodies when the incubation time with OTS964 was lengthened (**Fig. 4f**). This result suggests that OTS964 at low concentrations may partially inhibit CDK11 and, especially, even less effectively inhibit CDK10, leading to an incomplete shut down of the activity of CDK12. Another interesting observation is that there is an accumulation of Ser2ph-CTD of Pol II along gene bodies and at the transcription terminal sites (TTSs) (**Fig. 4f**). This becomes more obvious for a short period of inhibition of low concentration of OTS964 (**Fig. S5, S6**). The results suggest that CDK11 may be involved in the regulation of transcription termination besides elongation and splicing. It will be of great interest to investigate the cognate targets of CDK11 in the termination complex.

## Discussion

CDK11 was found to be essential for the embryonic development of animals two decades ago (Li, Inoue et al. 2004). However, the exact function and its cognate substrates remain largely unknown, though a recent report showed that the splicing factor SF3B1 is the substrate of CDK11 (Hluchy, Gajduskova et al. 2022). Here, we showed that CDK11 is the animal ortholog of Bur1 kinase from yeast and proved that both the CRT region of hSpt5 and the linker region between the hepatic repeats and the body of Rpb1 of Pol II are substrates of CDK11.

Furthermore, we found that CDK11 phosphorylates and activates CDK12 and, quite possibly, CDK13 as well. Moreover, we found that CDK10 is a partial homolog of CDK11, which also works on the linker region of Rpb1 and CDK12, but may not on hSpt5. Based on these discoveries, we propose that CDK11/10 is required to build up an elongation complex of Pol II starting from an initiated Pol II complex (**Fig. 5a**). First, the phosphorylation of CTR of Spt5 by CDK11 is required to recruit Paf1c (**Fig. 5b**). Furthermore, the phosphorylation of the linker region of Rpb1 by either CDK11 or CDK10 is required to recruit Spt6 (**Fig. 5c**). Moreover, the phosphorylation of CDK12 or CDK13 by CDK11 and CDK10 is essential for the generation of elongation active Pol II (Ser2ph-CTD of Pol II, **Fig. 5d**). Finally, CDK11 and/or CDK10 will couple with the elongating Pol II complex (**Fig. 5e**) to phosphorylate splicing factors and termination factors along the gene bodies (not shown).

**Figure 5.**
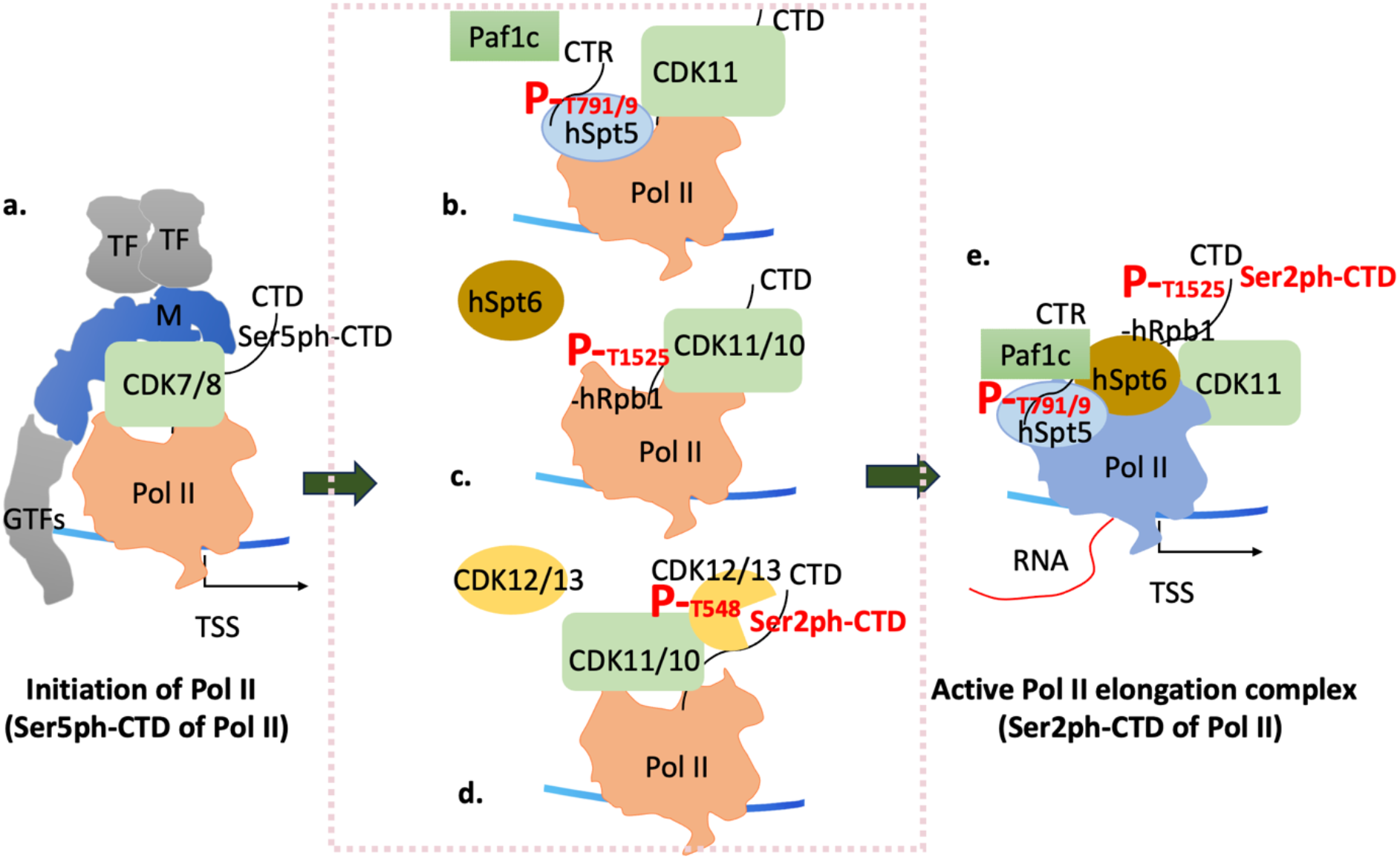
CDK11/10 phosphorylates CTR of hSpt5, the linker of Rpb1, and CDK12 to trigger the elongation of Pol II. **A**. T791/9-hSpt5 phosphorylation to recruit Pafic. **B**. Phosphorylation of the linker of Rpb1 to recruit hSpt6. **C**. CDK11/10 phosphorylates and activates CDK12/13, which phosphorylates Ser2-CTD of Pol II in turn. **D**. The building of elongation complex of Pol II by CDK11/10.

An interesting observation is that the phosphorylation site (T548, **Fig. 4d**) within CDK12 by CDK11/10 is located nowhere near the catalytic core but within the intrinsic disordered region (IDR) of CDK12. It is well-established that IDRs of proteins have the autonomous capacity to form condensates and phase separation from the hydrophilic surroundings. From sequence analysis, there are 10 “-TP-” sequences within human CDK12 and 8 “-TP-” sequences within human CDK13. One likely explanation is that the phosphorylation of these sites within the IDR by CDK11/10 disrupts the phase separation to expose the catalytic core of CDK12 and CDK13 for substrate recognition. Here, the Ser2-CTD of hepatic repeats of Rpb1 of Pol II is the cognate substrate of CDK12/13.

CDK9 has been thought, for decades, to be the animal ortholog of yeast Bur1. This has led to confusion about the underlying mechanism of promoter-proximal Pol II pausing, which only happens in metazoans as elegantly demonstrated by Lis and colleagues (Jonkers and Lis 2015). An early report from Eick’s group showed that CDK9 generates a unique phosphorylation pattern of CTD of Pol II in animals (Schuller, Forne et al. 2016), which is different from that of yeast (Schuller, Forne et al. 2016, Suh, Ficarro et al. 2016). Our group further characterized this unique phosphorylation pattern, which is only recognized by an antibody targeting consecutive repeat phosphorylated Ser2-CTD (Ser2ph-Ser2ph-Ser5ph-CTD) of Pol II but not by 3E10 and specifically generated by CDK9 (Liu, Ramachandran et al. 2020). We also found that this unique pattern is required to recruit JMJD5 via a C-terminal interaction domain (CID) (Liu, Ramachandran et al. 2020), responsible for cleaving arginine methylated histone tails of the +1 nucleosome in mice to generate fragile “tailless nucleosome” and the final release of paused Pol II into productive elongation (Liu, Wang et al. 2017, Liu, Wang et al. 2018), which is otherwise halted by the +1 nucleosome as demonstrated by four groups (Mavrich, Jiang et al. 2008, Schones, Cui et al. 2008, Gilchrist, Dos Santos et al. 2010, Weber, Ramachandran et al. 2014). Based on these discoveries from others and ourselves, we concluded that CDK9 is essential to trigger the release of promoter-proximal paused Pol IIs and only exists in higher eukaryotes (Lee, Liu et al. 2020, Liu, Ramachandran et al. 2020, Zhang 2024). Our data also showed that the unique phosphorylated pattern of CTD pulled down by JMJD5 is recognized by the 3E8 antibody, which is specific for Ser5ph-CTD of Pol II (Liu, Ramachandran et al. 2020). It suggests that CDK9 may work on initiated Pol II with the phosphorylated Ser5-CTD of Pol II, which is generated by CDK7 and CDK8, consistent with the early observation that initiated Pol II (Ser5ph-CTD of Pol II) is paused upon release after phosphorylation by CDK9 (Marshall, Peng et al. 1996, Lis, Mason et al. 2000). Based on these observations, we conclude that CDK9 works after CDK7 and CDK8 but before CDK12/13 (**Fig. 6**). From this report, we find that CDK12 is a cognate substrate of CDK10 and CDK11. It should be logical to put CDK10/11 or Bur1 ahead of CDK12/13 or Ctk1 (**Fig. 6**). Finally, a new working flow of cyclin-dependent kinases, which involves the transcription regulation, is established (**Fig. 6**). Since CDK9 functions similarly to those of CDK12 and CDK13 to generate Ser2ph-CTD of Pol II, it will be of great interest to figure out if the unique phosphorylation pattern is capable of recruiting splicing complex and termination complex without further action of either CDK11/10 or CDK12/13 for these Pol II pausing regulated genes. In this regard, CDK9 may directly bypass CDK11/10 and CDK12/13 (**Fig. 6**).

**Figure 6.**
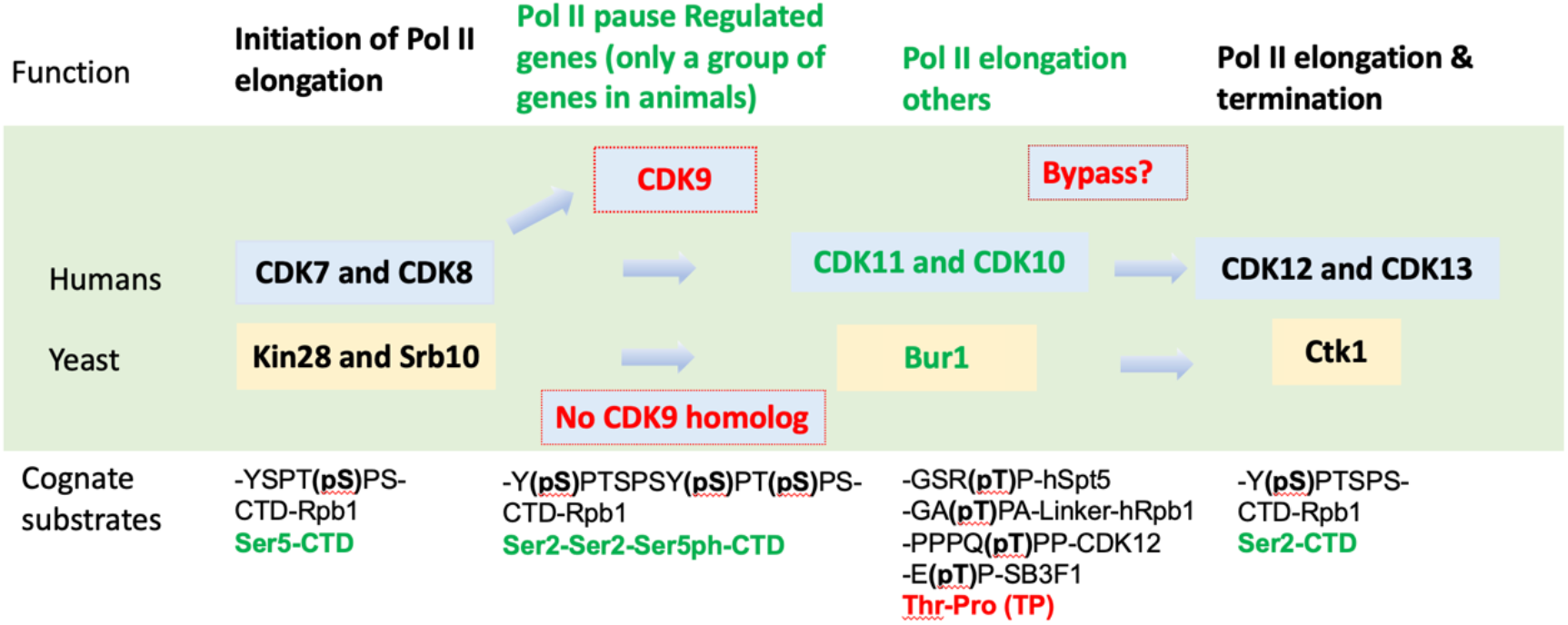
The potential stepwise function of major cyclin-dependent kinases in transcription regulation in eukaryotes. CDK9 is unique and only exists in metazoans, CDK7, CDK8, CDK10, CDK11, and CK12/CDK13 are conserved from yeast to humans. CDK10/11 is specific for **“Thr-Pro/TP”** signature sites while others prefer **“Ser-Pro/SP”**.

Our discovery that CDK11/10, instead of CDK9, is the ortholog of Bur1 kinase finally resolves this longstanding mystery. Besides hSpt5, the linker region of Rpb1, and splicing factor SF3B1 (Hluchy, Gajduskova et al. 2022), CDK12 is one of the cognate substrates of CDK11.

Unexpectedly, we found that CDK10 could partially compensate for the function of CDK11 and become a redundant homolog of CDK11 in metazoans. We expect that CDK13 could be a substrate of CDK11, too, since CDK13 is a highly similar homolog of CDK12 with exactly the same catalytic function as CDK12. Furthermore, we believe that some transcription termination factors could be targets of CDK10/11 since Bur1 also couples with elongating Pol IIs in yeast, as reported by Cramer’s group (Lidschreiber, Leike et al. 2013). As mentioned above, there are multiple similar “-TP-” sequences within both CDK12 and CDK13. It is astonishing that all substrates of CDK11/10 characterized have a signature sequence of “-TP-” as sites observed from hSpt5, the linker of Rpb1, SB3f1, and CDK12, which is different from “-SP-” of other transcription related cyclin-dependent kinases (**Fig.6**). This observation will help us to characterize unidentified substrates of CDK10/11.

## Methods

### CDK11 recombinant protein and antibodies

The recombinant human CDK11:1-502, Cyclin C: 1-283 protein is purchased from Thermo (A33511). The rabbit polyclonal CDK11 antibody is from Fortis Life Science (A300-311A). Rabbit phosphor-SPT5 polyclonal antibody is provided by FabGennix International Inc. Total SPT5 antibody is from Santa Cruz Biotechnology (sc-133217). Pol II CTD phospho-serine 2 (61083, 3E10) and phosphor-serine 5 (61085, 3E8) antibodies are from Active Motif. The Total Pol II antibody is from Santa Cruz Biotechnology (sc-56767). Pol II RPB1 phosphothreonine 1471 antibody is a gift from Drs. Tim Formosa and Christopher Hill’s groups. pT548-CDK12 antibody is a gift from Dr. Philippe Roux’s group.

### CDK11 shRNA knockdown in 293T cells

Human CDK11 shRNA plasmids are purchased from Sigma. Half a million 293T cells are seeded at the 6-well plate one day before transfection. 2 µg control or CDK11 shRNA is transfected into cells with Lipofectamine 2000. 48 hours after transfection, cells are harvested and lyzed with RIPA buffer. Equal amount of total cell lysate is loaded and separated by SDS PAGE gel, transferred to the PVDF membrane, and detected by the corresponding antibodies. For experiments of shRNA knockdown coupled with inhibition of CDK10 by Dinaciclib, 1μM Dinaciclib (MCE, #HY-10492) was added to the shRNA-treated cells and incubated for 10 minutes followed by cells harvest and lysis with RIPA buffer.

### OTS964 treatment of 293T cells

OTS964 is purchased from MedChemExpress. 800 thousand 293 T cells are seeded at the 6-well plate one day before treatment. OTS964 was incubated at the indicated concentrations for 1 hour. Cells were then lysed with RIPA buffer. The lysate was then analyzed by western blotting.

### MBP-spt5 or MBP-Pol II linker protein expression

The gene sequence encoding amino acid of human Spt5 from 754 to 811 or human Pol II RPB1 subunit 1511 to 1560 is cloned into a pMAL vector. The plasmid is transformed into BL21(DE3) competent E.coli cells. The MBP-spt5 or Pol II linker fusion protein is purified with amylose resin.

### CDK11 ^32^P-ATP radioactive kinase assay

**0**.1 µg CDK11 is incubated with 10 µg MBP-spt5 or MBP-Pol II linker fusion protein with 5 µCi ^32^P-ATP/sample in kinase buffer (50 mM HEPES, pH 7.5, 1 mM DTT, 2 mM MnCl_2_, 2 mM sodium orthovanadate). For the inhibition assay, 100 nM OTS964 is present in the reaction. The kinase reaction is carried out at 30 oC for 1h and stopped by adding SDS loading buffer and heating at 95 for 5 minutes. It’s then separated on the SDS PAGE and the gel is dried for 2 hours with heating at 80 oC and vacuum pumping. The radioactive signal is detected by the X-ray film.

### CUT&Tag-seq

CUT&Tag-seq was performed according to the published method (Kaya-Okur, Wu et al. 2019). Briefly, ∼100,000 HEK293T cells that were untreated, and treated with 200 nM or 1μM OTS964 for 2.5 minutes, 5 minutes, 7.5 minutes, 10 minutes, or 20 minutes were spun down and washed once with wash buffer. The cells were resuspended in 1.5 mL wash buffer and then bound to Concanavalin A-coated beads. The cells with beads were resuspended in 800 µL ice-cold antibody buffer containing RNA pol II CTD Ser2ph antibody from active Motif (#61083), then placed on a nutator at 4 °C and incubated overnight. After washing, resuspended the cells with beads in Dig-wash buffer with secondary antibody from Antibodies-online (ABIN102153, and placed the tubes on a nutator at room temperature for 30–60 min.

Then washed with 0.8 - 1 mL Dig-wash buffer three times. Next, the cells with beads were resuspended in Dig-300 buffer mix with pA-Tn5 adapter complex (C01070001, Diagenode, NJ). Place the tubes on a nutator at room temperature for 1 hr. After tagmentation, the DNA was extracted. Purified DNA was amplified using the following conditions: Cycle 1: 72 °C for 5 min; Cycle 2: 98 °C for 30 sec; Cycle 3: 98 °C for 10 sec; Cycle 4: 63 °C for 10 sec; Repeat Cycles 3-4 13 times; 72°C for 1 min and hold at 4 °C. The amplified libraries were purified, and size-selected, and the quality and quantity of libraries were assessed on 4150 TapeStation System (Agilent, CA). The pair-ended sequencing of DNA libraries was performed on an Illumina NovaSEQ6000 platform.

### CUT&Tag-seq data analysis

Raw sequencing reads were trimmed to remove adaptors and low-quality sequences using TrimmomaticPE (version 0.33). The trimmed sequencing reads were aligned to the hg38 reference genome using Bowtie2 (version 2.3.4.1) with very-sensitive and -x 2000 parameters. The read alignments were filtered using SAMtools (version 1.7) to remove PCR duplicates. Narrow peaks were called by MACS2 (version 2.1.2) with the q-value cut-off of 0.05. To analyze loss binding sites after OTS964 treatment, CUT&Tag-seq reads of HEK293T samples that were untreated were compared with those of HEK293T samples treated with OTS964 using the R package DiffBind (version 3.12.0) (Ross-Innes, Stark et al. 2012). All raw data were processed by Basepair (https://app.basepairtech.com/). Bam files from Basepair were used in DeepTools of Galaxy (https://usegalaxy.eu/) to generate bamCompare files, computeMatricx files, and final heatmap figures (Ramirez, Ryan et al. 2016).

## Acknowledgments

We thank Dr. Philippa Marrack for the final editing and suggestions. Thank Dr. Tony Gerber, and other members of National Jewish Health for their suggestions and support. Thank Drs.

Christopher Hill and Tim Formosa for antibodies against the phosphorylated linker of Rpb1 of yeast. The work is partially supported by an NIH grant GM135421 (G.Z.) and funds from NB Life Laboratory LLC, specifically private funds from Cheng-Yuan Zhang, Shi-Ning Xu, Lian-Hua Jin, Peng Sun, Yun-Xia Jiang, and Yongmei Jiang.

## Contributions

GZ conceived the concepts, designed experiments, conducted data analysis, and wrote up the manuscript. HL, LL, and JG carried out most experiments. TH and PR for the pT548-CDK12 antibody and suggestions and editing. SL, PW, QZ, and HH help in the final data analysis.

## Supplementary data

