## Supplementary material for "CDK11 activates CDK12 to trigger the elongation of RNA Polymerase II": no

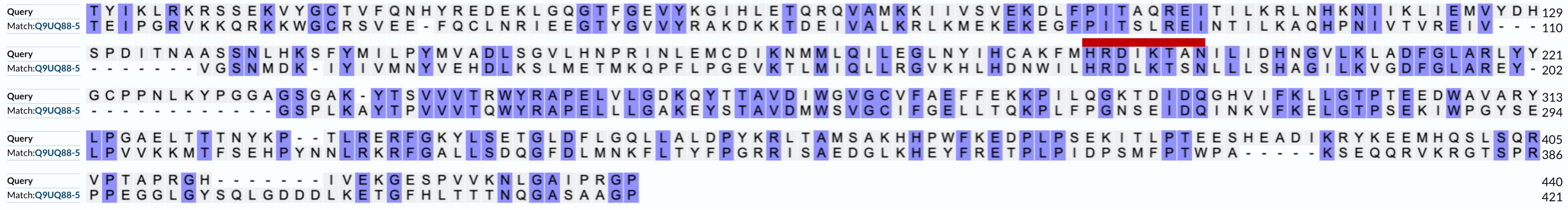

**Fig. S1.** The conserved "PITS(A)L(Q)RE" motif of CDK11 and Bur1. The BUR1 protein sequence from Saccharomyces cerevisiae (strain ATCC204508) is used for a BLAST search in the human genome. "PITSLRE " signature motif containing family members, CDK11A (Q9UQ88-5)/B (P21127-12) from the human genome pop up as the top two candidates with CDK9 after them.

| | |
|---|---|
| Human-hSpt5 | 772-**GSQTP**MYGS**GSRTP**MY**GSQTP**LQD**GSRTP**HY**GSQTP**LHD**GSRTP**-815 |
| S. pombe-Spt5 | 824-**GSRTP**AWNT**GSRTP**AWNS**GSKTP**AWNS**GSRTP**AWNS**GNKTP**AWNA**GSRTP**-873 |
| S. cerevisiae-Spt5 | 931-**SSWGGA**STWGGQGNGGA**SAWGGA**GGGA**SAWGGQ**GTGAT**STWGGA**SAWGNK**SSWGGA**STWASGGESNGAM **STWGGT**GDRSAYGGA**STWGGN**NNNKSTRDGGASAWGNQDDGNR**SAWNNQ**GNKSNYGGN**STWGGH**-1063 |

**Fig. S2. The multiple repeats of Spt5 from humans and yeast**. Top: Human (O00267) hSpt5 repeat sequence from 772-815. Middle: Schizosaccharomyces pombe (O13936) Spt5 repeat sequence from 824-873. Bottom: Saccharomyces Cerevisiae (P27692) Spt5.

**a. Conserved linker region of Rbp1 between yeast and human beings**

```
S. cerevisiae  AA1561-EIEDGQDGGVT(1471)PYSNESGLVNADLDVKDELMFS(1493)PLVDSGSNDAMAG-AA1506 linker of Rpb1
       Humans  AA1515-PAMTPWNQGAT(1525)PAYGAWSPSVGSGMTPGAAGFS(1547)PSAASDASGFSPG-AA1560 Linker of Rpb1
                          *         *                            **         *
```

**b. shRNA knockdown of CDK11 and pT1525-Rpb1**

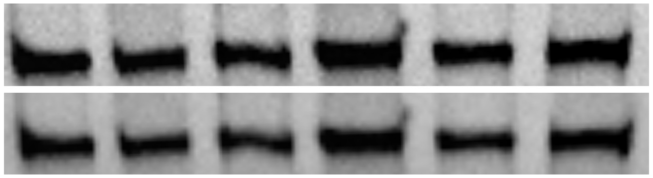


**Fig. S3. CDK11 specifically phosphorylates the linker-Rpb1 of Pol II. a**. sequence alignment of linker region of Rbp1 of yeast and human. **b.** shRNA knockdown of CDK11 does not affect the level of phosphorylated linker of Rpb1 in vivo.

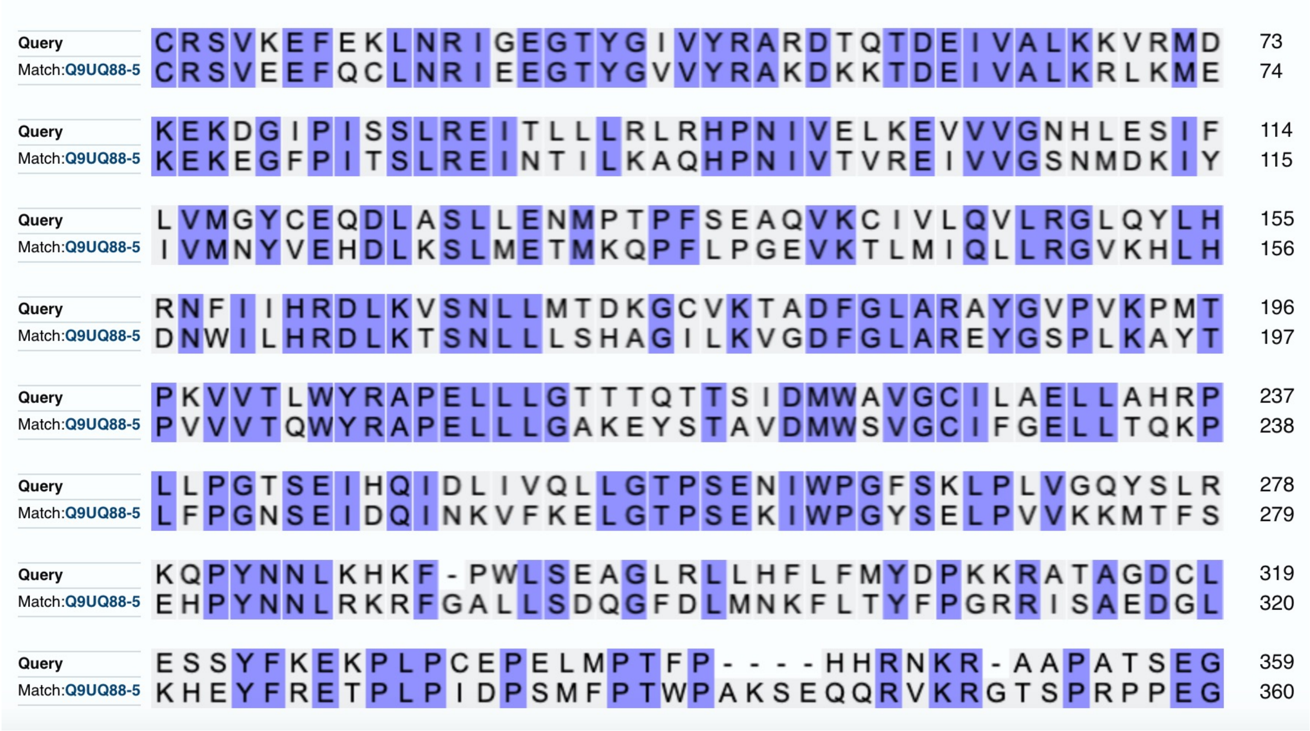


**Fig. S4. The sequence alignment of CDK10 and CDK11.** Top: The sequence of CDK10 from humans. Bottom: The sequence of CDK11 from humans.

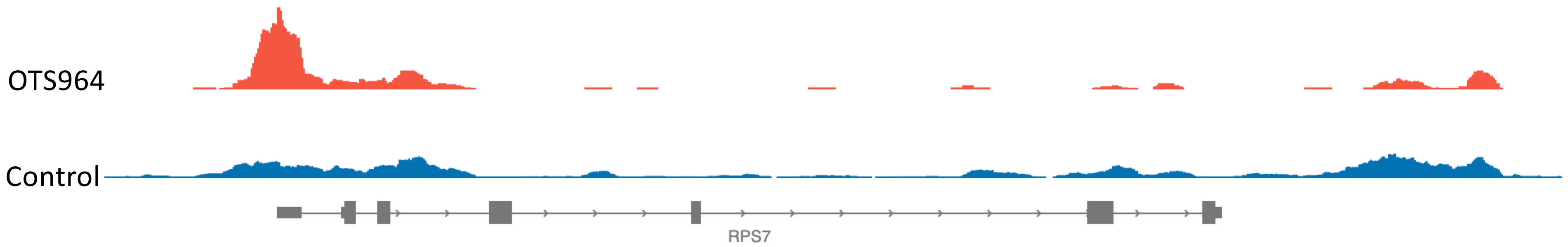


**Fig. S5.** Ser2ph-CTD of Pol II accumulates at the transcription start site with the inhibition of 200 nM OTS964.

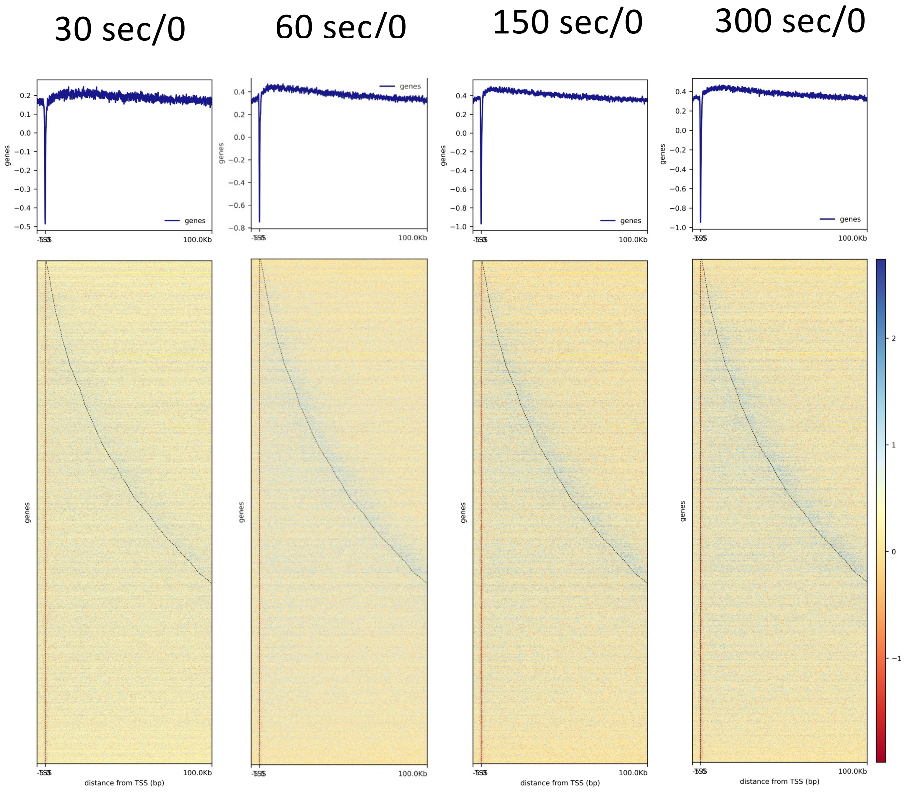


**Fig. S6.** OTS964 (1000 nM) inhibition with short time leads to accumulation of Ser2ph-CTD of Pol II along gene bodies and transcription termination sites (TTSs).

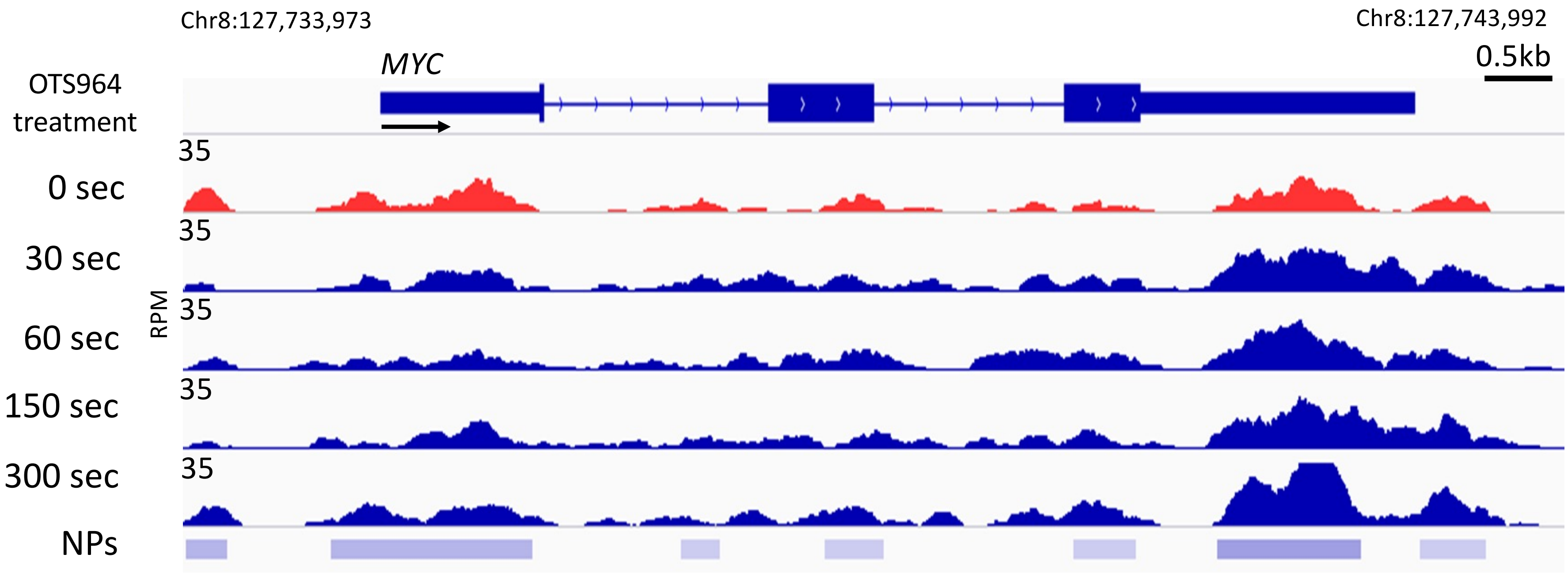


Fig. S7. Inhibition of OTS964 (1,000 nM) leads to the overall drop of active Pol II at transcription start sites (TSSs) but accumulates along gene bodies and at Transcription termination sites (TTSs). The numbers along the Y axis represent RPM: reads per $10^6$ mapped reads. The arrows indicate the direction of *MYC* gene transcription. "sec" = second. "NPs" = Group 0 sec CUT&Tag-seq narrow peaks.
